## Supporting Information for "Design of nanobody targeting SARS-CoV-2 spike glycoprotein using CDR-grafting assisted by molecular simulation and machine learning"

**S1 Script. Interface Analyzer from Rosetta**

**S1 Methods. Machine learning**

**S2 Methods. Molecular mechanics generalized Born surface area (MMGBSA)**

**S3 Methods. ELISA for the nanobody detection**

**S1 Fig. Illustration of the structure of the SARS-CoV-2 RBD-bound N-glycan attached to N- 343.**

**S2 Fig. Box plot for visually displaying the distribution and skewness of the interface parameters in 80 natural Nbs-antigen interfaces showing the data quartiles and averages.**

**S3 Fig. Accuracy of a K-Nearest Neighbors (KNN) model as the number of neighbors k increases from 1 to 10.**

**S4 Fig. Nanobody production and detection**

**S5 Fig. Production and morphological assessment of VLPs**

**S1 Table. Interface parameter for the filtered nanobodies from the total pool of designs**

**S2 Table. Summary of Molecular Mechanics Generalized-Born Surface Area (MM-GBSA) Calculations**.

### S1 Script. Interface Analyzer from Rosetta

```
<ROSETTASCRIPTS>
	<SCOREFXNS>
		<ScoreFunction name="ref2015" weights="ref2015"/>
	</SCOREFXNS>
	<FILTERS>
		<ShapeComplementarity name="Sc" min_sc="2.0" write_int_area="1" jump="1" 
confidence="0" />
		<Ddg name="ddg" scorefxn="ref2015" threshold="0" jump="1" repeats="5" repack="1" 
repack_bound="0" confidence="0" />
	</FILTERS>
	<MOVERS>
		<InterfaceAnalyzerMover name="ifa" scorefxn="ref2015" pack_separated="1" pack_input="1" 
tracer="0" interface_sc="1" interface="A_B"  />
	</MOVERS>
	<PROTOCOLS>
		<Add mover="ifa" />
		<Add filter="Sc" />
		<Add filter="ddg" />
	</PROTOCOLS>
</ROSETTASCRIPTS>
```

#### S1 Methods. Machine learning

The structures of nanobodies (Nbs) complexed with antigens, curated from the dataset available from Zavrtanik and Hadži [1], were geometry-relaxed using the FastRelax protocol in the Rosetta suite [2]. Coordinate constraints were applied to backbone heavy atoms based on the input structure, utilizing the following command:

**$ relax.macosclangrelease -ignore_unrecognized_res -relax:constrain_relax_to_start_coords -ex1 -ex2 -use_input_sc -s *.pdb**

Following relaxation, interface parameters were computed using the InterfaceAnalyzer mover from Rosetta. These computed descriptors served as input for feature selection, where an extra tree classifier with the default parameters of scikit-learn identified the 10 most relevant descriptors based on their importance. Each decision tree in the ensemble splits the data to minimize Gini impurity and maximize class separation. The decrease in impurity at each split reflects the feature's contribution to classification. Feature importance is the average impurity decrease across all trees for that feature.

These selected variables were crucial in predicting binding affinity, categorized as either "high" or "low" based on an energy threshold of -11.5 kcal/mol. Specifically, affinities were labeled "high" if the energy was below -11.5 and "low" otherwise. To prepare the data for model training, features were standardized using the StandardScaler from the scikit-learn library, ensuring a mean of 0 and a standard deviation of 1, followed by Linear Discriminant Analysis (LDA) to reduce dimensionality while maximizing class separability. The LDA-transformed data was split into training and testing sets in a 70:30 ratio, with stratification to maintain class proportions, using the train_test_split function with a random state of 42 for reproducibility. A K-Nearest Neighbors (KNN) classifier, with k=6, was trained on the LDA-transformed training data and used to predict class labels on the test set. Model performance was evaluated using accuracy, precision, recall, ROC AUC, and Matthews Correlation Coefficient (MCC). To validate the model's stability and generalizability, a 5-fold stratified cross-validation was performed, with the mean values of the evaluation metrics calculated across the folds to assess overall performance. The analysis was conducted using Python 3.8.10 with Pandas [3] for data management and scikit-learn [4] for data splitting, cross-validation, feature standardization, dimensionality reduction, KNN implementation, and performance evaluation.

**S2 Methods. Molecular mechanics generalized Born surface area (MMGBSA)**

The output from ClusPro [5] was used as the starting structure for MMGBSA simulations. The models were used directly from ClusPro, and therefore, without glycosylation. Initially, a 100 ns molecular dynamics simulation was performed, following the same setup as described in the main text. The resulting trajectories were then post-processed using the default parameters of the gmx_MMPBSA [6] pipeline. For the MMGBSA calculations, 400 frames were analyzed, utilizing the LCPO surface area method [7]. The calculations were conducted at a temperature of 303.15 K, with a saline concentration of 150 mM NaCl, and the modified GB model 2 developed by A. Onufriev, D. Bashford, and D.A. Case (GB-OBC2) [8].

**S3. Elisa for nanobody detection**

High binding, half area 96-well polystyrene plates (Costar; Lowell, MA, USA) were coated with 1 µg/mL of the Nb Ab.2 in 0.2 M carbonate/bicarbonate buffer (Pierce, IL, USA) overnight at 4 ºC. Plates were blocked with skimmed milk (Bio-Rad) at 5% (w/v) in PBS-T buffer [1X PBS with 0.05% (v/v) Tween 20] for 2 hours at room temperature. Anti-histidine monoclonal antibody (Rockland, USA) was diluted (1:1000) in an assay buffer [5% (w/v) skimmed milk in PBS-T] and added to the plate. After a 2 hours incubation at room temperature (RT), the plate was washed five times with PBS-T and incubated with horseradish peroxidase (HRP)-conjugated antibody against mouse IgG (Jackson ImmunoResearch, 1:20 000 dilution) diluted in assay buffer for 1 hour at RT. After a second round of washes, the reaction was developed by the addition of the enzyme substrate (tetramethylbenzidine TMB-KPL substrate (Pierce, IL, USA)) for 30 minutes at room temperature, followed by 1N HCl. Optical density at 450nm (OD450nm) was read in a microplate spectrophotometer (BioTek; Winooski, VT, USA). Graphpad Prism V.7 software (GraphPad Software, San Diego CA, USA) was used for statistical analysis.

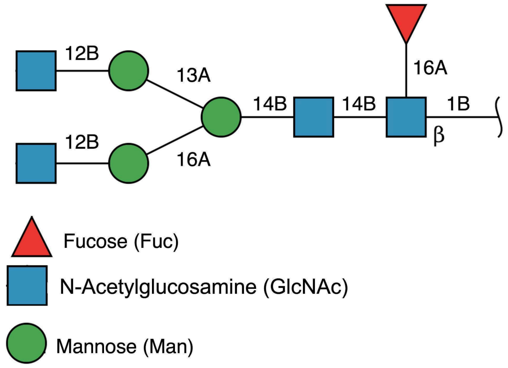


**S1 Figure. Illustration of the structure of the SARS-CoV-2 RBD-bound N-glycan attached to N- 343.** The N-glycan is composed of three components: N-Acetylgucosamine (depicted by a blue square), mannose (depicted by a green circle), and fucose (depicted by a red triangle). The $\alpha$ and $\beta$ linkage types are represented by A and B, respectively. The numbering between the glycosidic linkages indicates the carbon involved in the bond, where the first number is the carbon number of the first monosaccharide and the second number is the carbon number of the second monosaccharide.

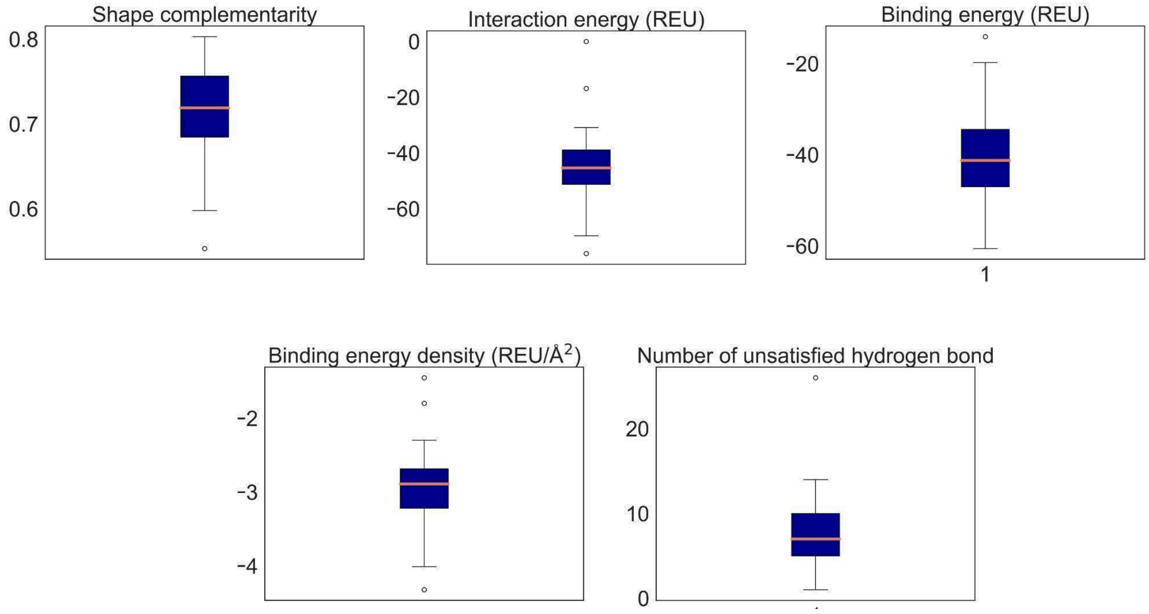


**S2 Figure. Box plot for visually displaying the distribution and skewness of the interface parameters in 80 natural Nbs-antigen interfaces showing the data quartiles and averages.** The interquartile range is shown as a solid blue box, where the top and bottom of the box denote the upper and lower quantile, respectively. The median is depicted as an orange line. Outliers are represented by circles. REU = Rosetta energy units.

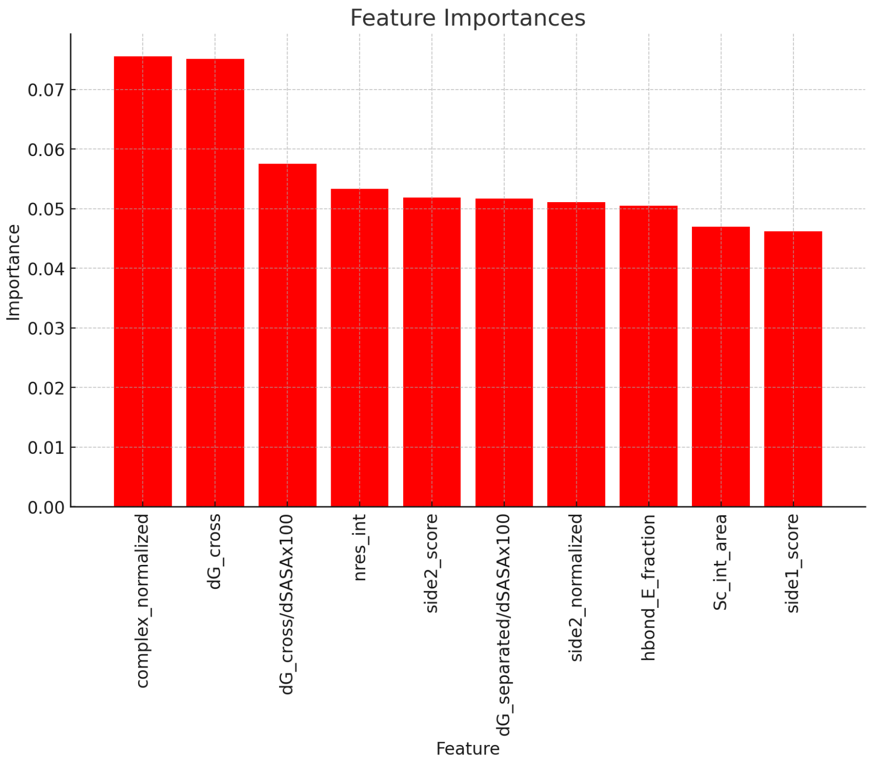


**S3 Figure.** The top 10 most important features based on the Extra Trees classifier.

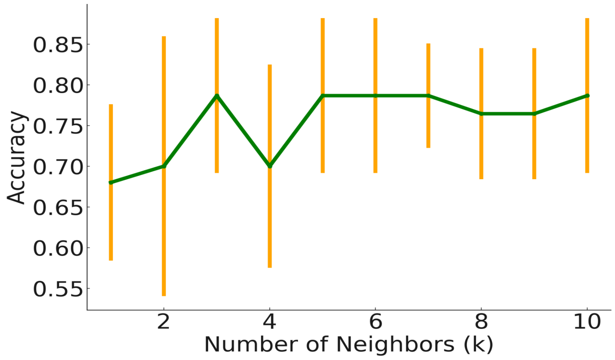


**S4. Accuracy of a K-Nearest Neighbors (KNN) model as the number of neighbors k increases from 1 to 10.** The green line represents the mean accuracy, while the orange bars indicate the standard deviation. Accuracy peaks at k=5, though variability is high, especially for lower k values.

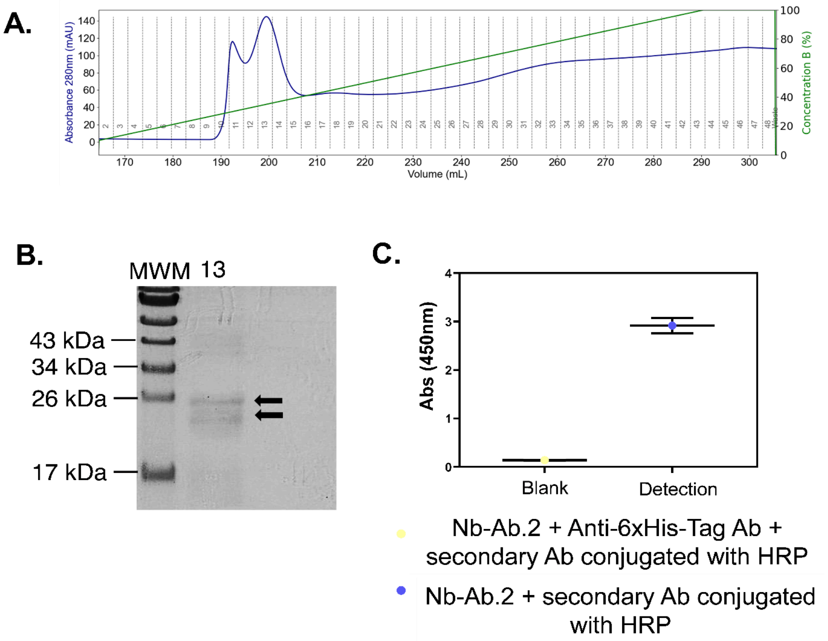


**S5 Figure. Nanobody production and detection. (A)** Chromatogram from IMAC Purification of Nb-Ab.2. The x-axis represents the elution volume (mL), while the left and right y-axes show the absorbance at 280 nm (A280) and the concentration of the elution buffer (%), respectively. **(B)** Fraction 13 was subjected to analysis using SDS-PAGE (17.5%). The nanobody Nb-Ab.2 (19.5 kDa) migrated within the polyacrylamide gel with the expected mass, around 20 kDa. The occurrence of an additional band on the gel, positioned above the target band, is likely indicative of an incomplete removal of the pelB signal peptide from the N-terminal sequence of the nanobody [9]. The yield obtained was 0.5 mg per liter of bacterial culture. **(C)** Nb detection was made by using an in-house His-Tag Protein ELISA assay. Plate was coated with the purified Nb overnight. The next day, an anti-6xHis monoclonal antibody was added, followed by an HRP conjugated secondary antibody. We used the signal generated solely by the addition of the secondary antibody to the plate as our blank control.

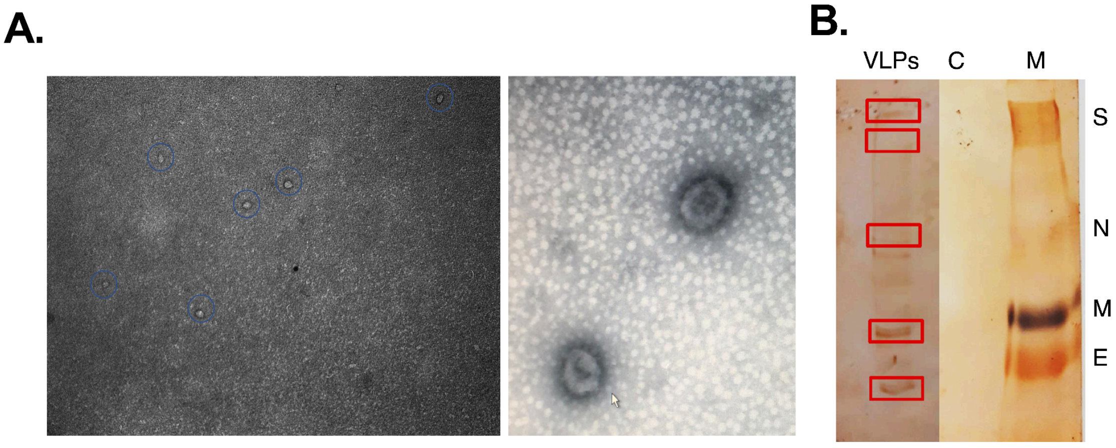

**S6 Figure. Production and morphological assessment of VLPs. (A)** Transmission electron microscopy **(**TEM) images of SARS-CoV-2 VLPs produced in Vero E6 cells 48 h after co-transfection with the plasmids encoding the virus structural proteins (M, S, E, and N) according to the 8:6:8:3 molar ratio. (B) Western-blot analysis of purified VLPs. Detection was performed using a pool of SARS-CoV-2-infected individuals.

**Table S1. Interface parameter for the filtered nanobodies from the total pool of designs.**

| Interface Parameter | Nb Ab.1 | Nb Ab.2 |
|---|---|---|
| Shape complementarity | 0.71 | 0.71 |
| Binding energy (REU) | -33.20 | -37.27 |
| Interaction energy (REU) | -41.23 | -45.77 |
| Binding energy density (REU/Å) | -3.10 | -3.17 |
| Unsatisfied H-bond | 7 | 6 |
| RMSD post-FastRelax | 0.90 | 0.95 |

**Table S2. Summary of Molecular Mechanics Generalized-Born Surface Area (MM-GBSA) Calculations**. The estimated free energies, in kcal/mol, were decomposed to evaluate the total binding free energy components for the interaction between the two most likely docking structures of the designed Nb Ab 2 and SARS-CoV-2 RBD based on cluster size (model 0) and cluster energy (model 7)

| | Model 0 | | Model 7 | |
|---|---|---|---|---|
| Energy component (kcal/mol) | Average | SD | Average | SD |
| ΔVDWAALS | -26.56 | 3.65 | -50.46 | 5.20 |
| ΔEEL | -110.85 | 28.50 | -190.15 | 45.55 |

| | | | | |
|---|---|---|---|---|
| Δ1-4 VDW | 0.00 | 0.00 | 0.00 | 0.00 |
| Δ1-4 EEL | 0.00 | 0.00 | 0.00 | 0.00 |
| ΔEGB | 129.86 | 26.21 | 213.46 | 39.28 |
| ΔESURF | -4.33 | 0.51 | 06.90 | 0.62 |
| ΔGGAS | -137.41 | 28.00 | -240.61 | 45.02 |
| ΔGSOLV | 125.53 | 26.04 | 206.55 | 38.99 |
| **ΔTotal** | **-11.88** | **3.48** | **-34.06** | **7.79** |
